# Cell cycle and developmental control of cortical excitability in *Xenopus laevis*

**DOI:** 10.1101/2022.02.11.480124

**Authors:** Zachary T Swider, Ani Michaud, Marcin Leda, Jennifer Landino, Andrew B. Goryachev, William M. Bement

## Abstract

Interest in cortical excitability – the ability of the cell cortex to generate traveling waves of protein activity – has grown considerably over the past twenty years. Attributing biological functions to cortical excitability requires an understanding of the natural behavior of excitable waves and the ability to accurately quantify wave properties. Here we have investigated and quantified the onset of cortical excitability in *X. laevis* eggs and embryos, and the changes in cortical excitability throughout early development. We found that cortical excitability begins to manifest shortly after egg activation. Further, we identified a close relationship between wave properties – such as wave frequency and amplitude – and cell cycle progression as well as cell size. Finally, we identified quantitative differences between cortical excitability in the cleavage furrow relative to non-furrow cortical excitability and showed that these wave regimes are mutually exclusive.

## Introduction

The cell cortex, broadly defined as the plasma membrane and its proximal cytoplasm, represents the intersection between the cell and its environment. It is typically rich in signaling and cytoskeletal proteins, which allow it to integrate chemical and mechanical cues from both intracellular and extracellular sources, and appropriately exert physical changes in response to these signals. These changes are often dynamic and easily observable, such as extending cell protrusions during migration or pinching the cell in two during cytokinesis. Consequently, the cell cortex has attracted the attention of biologists for well over a century. However, while many cortical behaviors have been thoroughly studied and are well understood, new and important cortical behaviors continue to emerge.

One such behavior is cortical excitability, loosely defined as the ability of the cell cortex to generate traveling waves of filamentous actin (F-actin) polymerization and F-actin regulator protein recruitment. Excitability and related dynamics, such as oscillations, are thought to result from intertwined positive and negative feedback loops, with the former driving the traveling wave front forward at a constant velocity and the latter following closely behind to extinguish the trailing edge of the wave (for review see Michaud *et al*., 2021). While cortical waves of F-actin were first reported over twenty years ago (Vicker, 2000), improvements in imaging technology and molecular probes have led to a steady increase in the number of cell types reported to exhibit cortical excitability. Cortical excitability appears to be particularly common in locomoting cells, but it has recently been detected in distinctly non-motile cell types including those of the germline (Chanet and Huynh, 2020), stationary adherent cells (Graessl *et al*., 2017), and early embryos (Bement *et al*., 2015; Maître *et al*., 2015; Michaux *et al*., 2018), suggesting that the ability to generate waves may be a general feature of the cell cortex. The functions ascribed to cortical excitability are as diverse as the cell types within which it has been described and include establishing new adhesions (Case and Waterman, 2011), facilitating cell spreading (Hui *et al*., 2014), facilitating cell locomotion (Weiner *et al*., 2007; Miao et al., 2019) controlling the mode of cell locomotion (Stankevicins *et al*., 2020), driving cell compaction (Maître *et al*., 2015), regulating spindle orientation (Xiao *et al*., 2017), and promoting cytokinesis (Bement *et al*., 2015).

Cortical excitability remains a relatively new and sparsely studied process and, as such, a number of outstanding questions about it have yet to be answered. For example, what happens during the transition from a quiescent cell cortex to an excitable one? What is the relationship between cortical excitability and the cell cycle? Do waves scale with cell size? Does cortical excitability manifest differently in different parts of the cell cortex? While the answers may depend on the cell type and the behavior that cortical excitability is associated with, it is nonetheless of broad interest to follow and quantify features of natural cortical excitability in a single system as it changes and proceeds through different physiological states.

Here we have exploited the early development of *Xenopus laevis* to study cortical excitability. In many ways, this system is ideally situated to answer the questions posed above: The rapid rate of cell division in *X. laevis* embryos and concurrent reduction in cell size makes this system well suited to address how these factors influence cortical excitability. Moreover, the early development and cell cycle regulation of these embryos are well understood. Finally, cortical excitability in this system is constitutive and relatively easy to monitor, which allows for accessible imaging and extraction of quantitative metrics.

## Results

### Egg activation initiates cortical excitability

Cortical excitability is absent in immature *X. laevis* oocytes, but clearly evident in developing embryos and in artificially activated eggs several hours after activation (Bement *et al*., 2015). Similar oscillatory dynamics have also been reported using synthetic cortices based on supported lipid bilayers and activated egg cytoplasm (Landino et al., 2021). However, when and how the transition to cortical excitability occurs remains unknown. To determine this, we exploited the fact that traveling F-actin waves – the manifestation of cortical excitability – can be clearly detected in cells expressing the F-actin probe UtrCH (Burkel *et al*. 2007, Figure 1A, A’; Video 1). Moreover, UtrCH-labeled samples also allow quantification of informative wave metrics, namely, wave period (time between consecutive waves), temporal width, and peak-to-trough wave amplitude, all of which can be determined by following UtrCH fluorescence over time (Supplemental Figure 1A). As reported previously, these measurements can be made with sub-frame accuracy by simultaneously measuring n sub-regions of a field of view and averaging the results (Landino et al. 2021; Supplemental Figure 1B). In the experiments and analyses below, we have incorporated this approach into an analysis pipeline using the free and open-source language Python 3, and further expanded it to allow analysis of changing wave properties over time (Supplemental Figure 1C).

**Figure 1.**
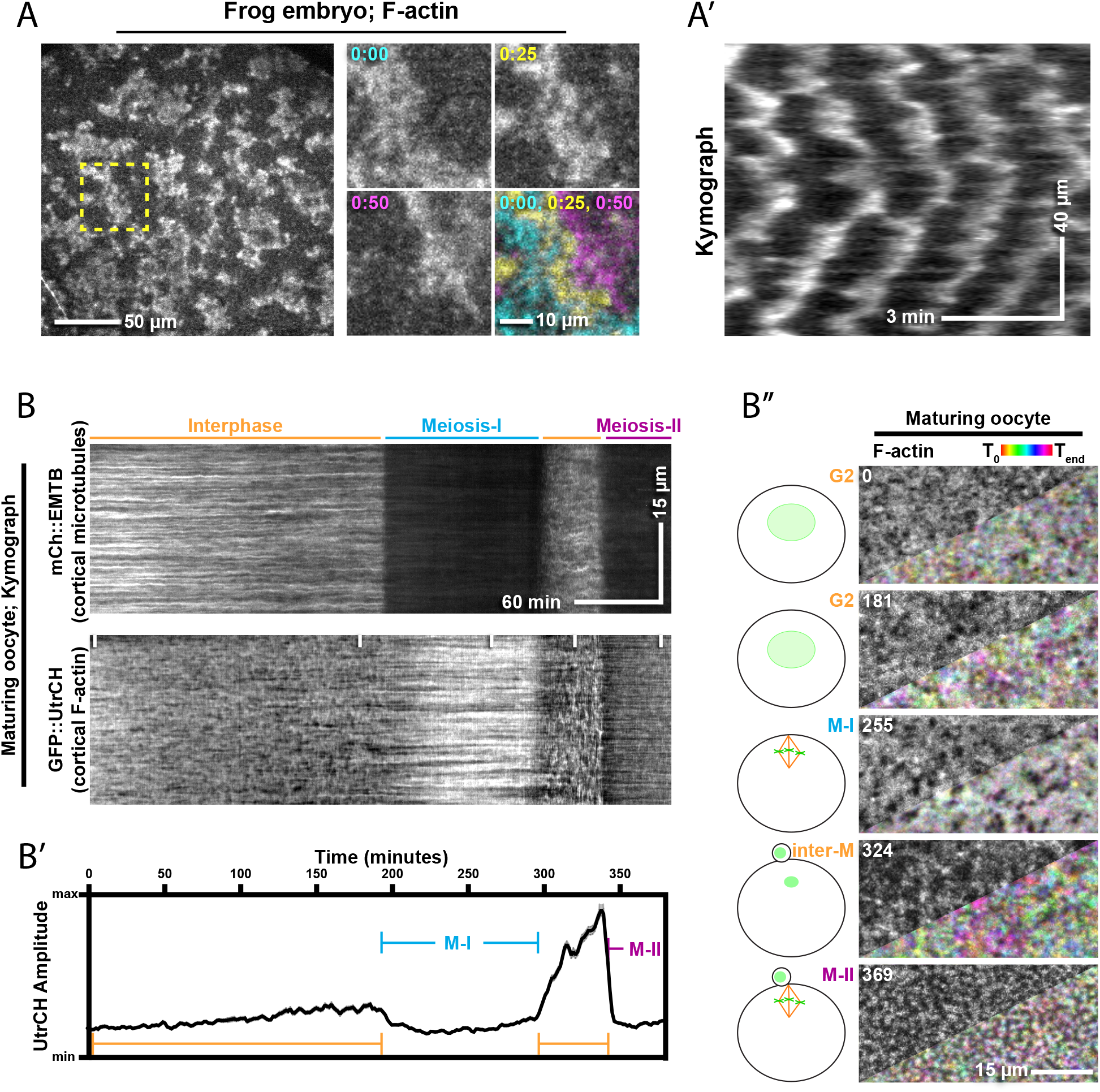
Analysis of excitable dynamics in maturing oocytes. (A) Still frame from an interphase ∼stage 5 embryo expressing a probe for F-actin (GFP-UtrCH). Three con-secutive times points at 25 second intervals are shown from the region outlined in yellow. A color-coded merge of the three time points is shown to illustrate the forward movement of the UtrCH labeling. (A’) Kymograph drawn from time-lapse used to create A illustrating the periodic nature of the waves in space over time. (B) Kymograph from a progesterone-treated oocyte undergoing maturation. The oocyte is expressing mCh-EMTB (microtubules; top kymograph) and GFP-UtrCH (F-actin;bottom kymo-graph). (B’) Plot showing changes in UtrCH amplitude (signal max-min over a 14 minute window) over time. The plot is positioned to match the kymograph above, and relevant cell cycle events (interphase, meiosis-I, and meiosis-II) are labeled consistent with the kymographs. (B’’) Regions marked with vertical lines on the GFP-UtrCH kymograph in B showing still frames in grayscale and a color-coded merge of a 1000 second time frame. Regions with mostly unchanging signal show up as gray while regions with changing signal are colored. Schematic diagrams illustrate cell cycle state. Time in minutes is shown in the top right of each still.

To identify the developmental transition to cortical excitability, we first monitored cortical F-actin dynamics during the process of meiotic maturation by treating immature oocytes with progesterone. Due to the large size of *Xenopus* oocytes (∼1.2mm in diameter), the germinal vesicle (the large oocyte nucleus) is too deep in the cytoplasm to use canonical markers for cell cycle state, such as DNA condensation. Thus, to simultaneously monitor meiotic progression and cortical F-actin dynamics, oocytes were microinjected with fluorescent probes for F-actin (UtrCH) and microtubules (EMTB; Faire *et al*., 1999). EMTB distinguishes between interphase and M-phase in that cortical microtubules are present during interphase but withdraw from the cortex during M-phase (von Dassow *et al*., 2009; Supplemental Figure 2A, A’). Indeed, EMTB labeling was consistent with the expected progression through meiosis; a long stretch of interphase with abundant cortical microtubules followed by meiosis-I where cortical microtubules were lost, a brief interphase-like state where cortical microtubules were once again abundant, and finally a prolonged arrest in meiosis-II where cortical microtubules were again lost (Figure 1B; Video 2). Despite prolonged imaging with high temporal resolution, we were unable to detect the presence of a periodic signal in either immature or maturing oocytes. However, we identified a reproducible increase in F-actin dynamics, defined here as the difference between signal minimum and maximum over time, as oocytes approached entry into meiosis-I. Dynamic actin once again returned to low levels during meiosis-I (Figure 1B-1B’’). Similar to previous reports in Echinoderm oocytes (Bement *et al*. 2015 and Bischof *et al*. 2017), we detected wave-like oscillations of F-actin and a large increase in dynamic actin following exit from meiosis-I (Figure 1B-1B’’; Video 2). We found these features to be suggestive of excitable dynamics; however, unambiguous waves were difficult to pinpoint and were not uniform across the field of view. Finally, as matured oocytes arrested in metaphase of meiosis-II, the cortex returned to a quiescent state with sharply reduced F-actin dynamics (Figure 1B-1B’’).

During the natural course of development, a mature (i.e., metaphase-II arrested) oocyte would be laid by the frog and fertilized externally. While *in vitro* fertilization is routine, imaging the events preceding and following fertilization is impractical as it requires an intact jelly coat which both hinders microinjection and makes high resolution imaging impossible. Accordingly, we activated mature (meiosis II arrested) oocytes by treating them with the calcium ionophore ionomycin, a widely used approach to mimic fertilization by elevating intracellular free Ca^++^. Ionomycin treatment evoked dramatic F-actin recruitment and simultaneous contractions of the cell cortex (seen in Figure 2B), consistent with the normal response to fertilization (Hara and Tydeman, 1979). Once cortical contractions stabilized ∼15 minutes after ionomycin exposure, cortical waves of F-actin with a period of 58±6 seconds (mean ± sd n = 3 cells 15 minutes after activation) could be detected in kymographs and fluorescence traces over time (Figure 2A-A’’; Video 3). To confirm that these findings were consistent with sustained cortical excitability, we performed extended time lapse imaging with sufficient temporal resolution to capture wave dynamics. Cortical waves were indeed sustained, but not uniform over time (Figure 2B; Video 4). Quantification of wave amplitude and wave period revealed that the normalized wave amplitude steadily increased over time, while the wave period initially decreased and then subsequently began to increase at about 20 minutes post-washout of ionomycin (Figure 2B’).

**Figure 2.**
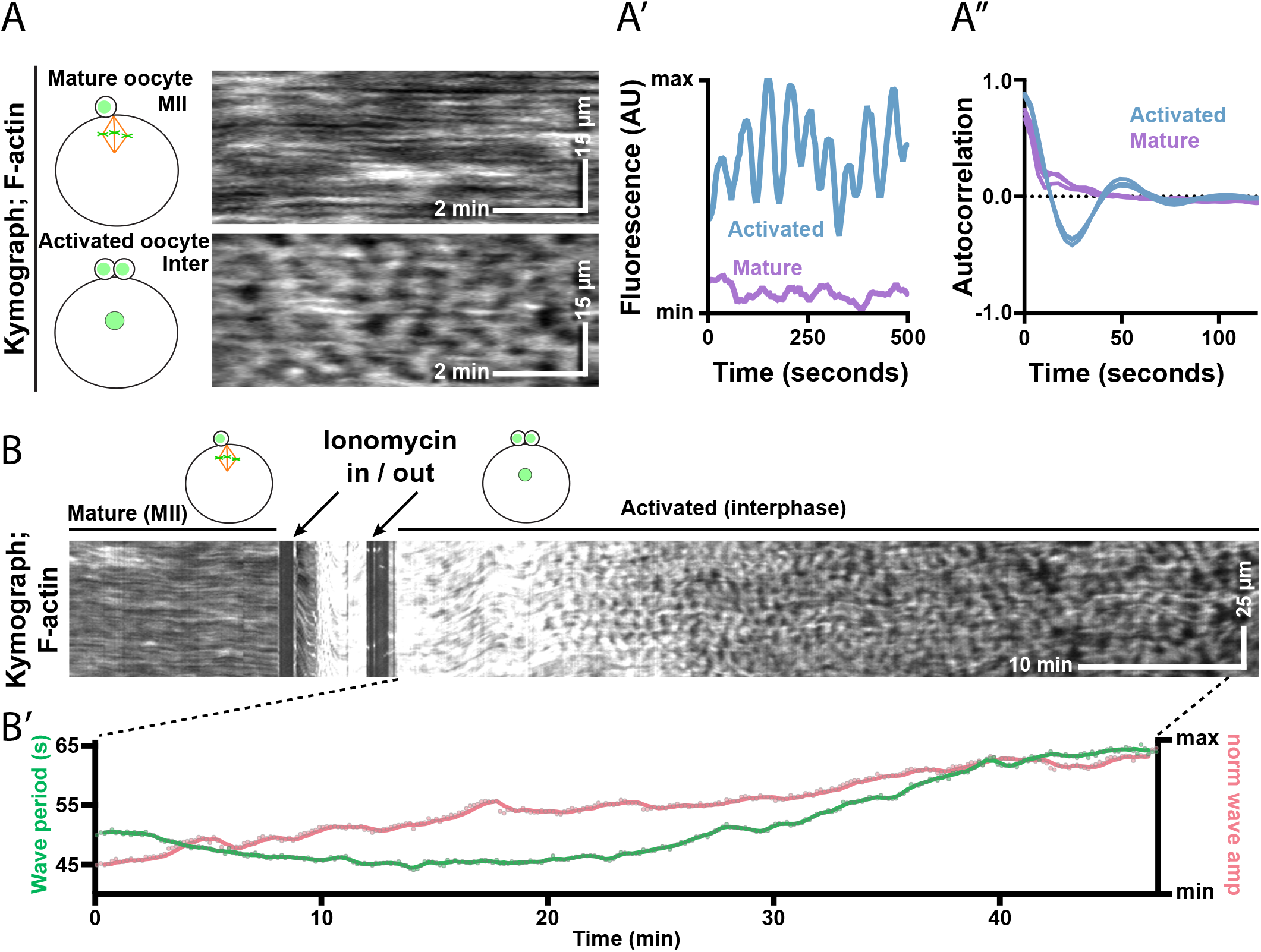
Excitability is detected after egg activation. (A) Kymographs from a mature (top) and an oocyte about 15 minutes following activation (bottom) expressing GFP-UtrCH (F-actin). Schematic diagrams illustrate cell cycle state. (A’) Representative trace from a ∼8μm2 ROI from the oocytes shown in A illustrating the presence of high-amplitude periodic waves in the activated but not the mature oocyte. (A’’) Mean autocorrelations from the mature and activated oocytes shown in A illustrating the presence of uniform periodic structures in the activated, but not the mature oocyte (n = three mature and three activated oocytes). (B) Kymograph from a GFP-UtrCH expressing oocyte responding to ionomycin treatment. This cell was treated differently and thus was not included in the analysis presented in A. Arrows on the kymograph indicate where ionomycin was flowed onto and washed off of the cell. (B) Plot showing the period and normalized wave amplitude for the region indicated in B.

### Wave properties oscillate with the cell cycle

Our previous analysis of wave dynamics in *X. laevis* revealed only slight changes over the cell cycle (Bement *et al*., 2015). However, this analysis included limited developmental stages, and was largely qualitative. Thus, with improved imaging and quantitative analysis, we sought to determine whether the changes we observed throughout the first interphase were representative of subsequent cell cycles. While the first mitotic interphase is relatively long (1-2 hours), subsequent cell cycles are comparatively rapid (15-30 minutes), allowing the recording of several cell cycles from a single cell (Hara *et al*., 1980).

Low magnification imaging of whole embryos suggested that the cortical cytoskeleton transitioned between loosely packed and densely packed waves prior to the appearance of a cleavage furrow (CF; Figure 3A; Supplemental Figure 3A, A’; Video 5). However, the normal cortical contractions associated with cytokinesis made long-term, high-resolution imaging difficult. To perform continuous high-resolution imaging, we instead imaged activated oocytes – which do not undergo cytokinesis, but otherwise exhibit normal mitotic biochemistry (Gerhart *et al*., 1984) – and, as described above, exploited the presence or absence of cortical microtubules to infer cell cycle state. Consistent with the imaging of embryos, we found that wave properties are in a continuous state of change in activated oocytes (Figure 3B; Video 6). The wave period consistently rose slowly throughout interphase, peaked prior to mitotic entry, and then “reset” back to early interphase characteristics beginning approximately halfway through mitosis (Figure 3B, B’). In contrast, wave amplitude peaked early in interphase and then remained steady or declined slightly as the cell approached mitosis. While wave peak and trough values were both consistently the lowest as the cells entered mitosis, the subsequent rise in wave trough values resulted in the lowest wave amplitude values about halfway through mitosis. Wave amplitude measurements rose rapidly as the cells exited mitosis, due to the continuing rise in wave peak and rapid drop in wave trough (Supplemental Figure 4).

**Figure 3.**
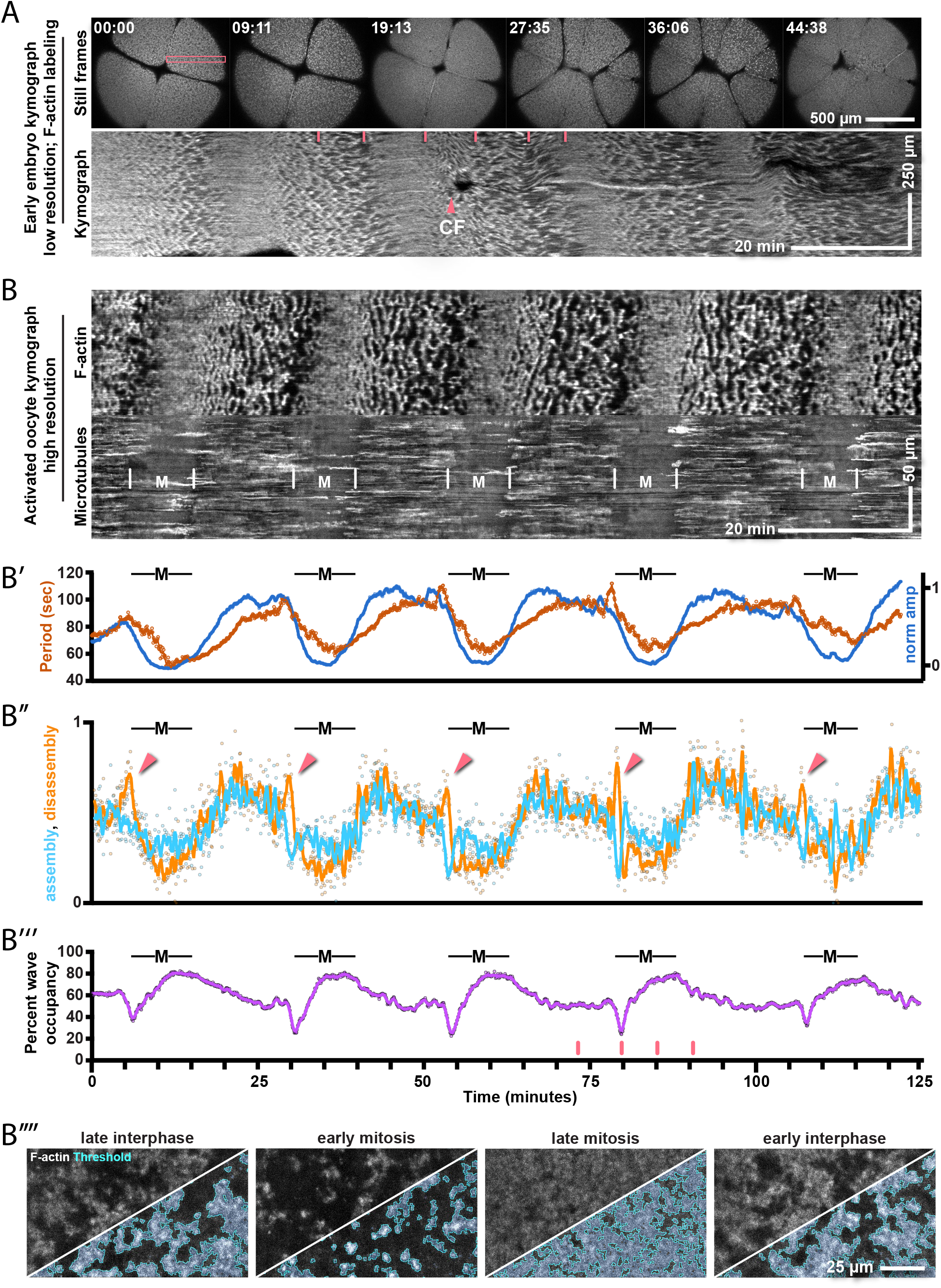
Cortical excitability fluctuates with cell cycle state. (A) Still frames (top) and kymograph (bottom) of a young embryo injected with Alexa647 conjugated UtrCH protein (F-actin). The kymograph was drawn from the region indicated in the pink box on the first still frame. The still frames correspond to the regions marked with vertical lines on the kymograph. The appearance of a cleavage furrow on the kymograph is marked with “CF”. (B) Kymograph from an activated oocyte expressing GFP-UtrCH (top kymograph) and mCh-EMTB (bottom kymograph). Entry into and out of mitosis, approximated from changes in EMTB labeling, are marked with vertical dashed lines. (B’) Plot showing wave period (orange) and normalized amplitude (blue) measurements over time, from cell shown in (B). Measurements were individually calculated from a 200 second window that rolled over the full dataset at 5 second intervals. (B’’) Plot showing total net F-actin assembly (light blue) and total net F-actin disassembly (light orange) over a 60 second window. Points on the plot represent single measurements, solid lines represent a rolling average of 5 frames. Pink arrows indicate regions where total net assembly and disassembly significantly diverge from one another. (B’’’) Plot showing the percent of the field of view occupied by F-actin waves at each time point. Points on the plot represent single measurements, solid lines represent a 25 second rolling average. Markings for mitotic entry and exit in B’-B’’’ are consistent with the markings in B. (B’’’’) Still frames from the kymograph in B corresponding to the time points indicated by the vertical pink dashes in B’’’. UtrCH fluorescence is shown in grayscale and thresholded wave regions are shownd outlined and highlighted in light blue.

These robust changes in wave behavior throughout mitosis reveal remodeling of the cortical F-actin cytoskeleton. Importantly, remodeling could result from a change in the rate of F-actin assembly, a change in the rate of F-actin disassembly, or both. To investigate how these factors shaped cortical waves during mitosis, we used UtrCH fluorescence as a readout for the quantity of F-actin present and measured the change in intensity of each pixel over the preceding n seconds as a readout for the relative rates of F-actin assembly and disassembly. Because the waves are insensitive to myosin-2 inhibition (Bement *et al*., 2015), and are thus not likely to reflect motor-mediated transport, we reasoned that if a pixel value increased over time that this reflected <u>net</u> F-actin assembly in that region and that if the pixel value decreased that this reflected <u>net</u> F-actin disassembly in that region. These measurements revealed that entry into mitosis is marked by a decrease in net F-actin assembly and a simultaneous increase in net F-actin disassembly (Figure 3B’’; Video 7). This change results in a sharp decrease in the percent of the cell cortex occupied by F-actin from ∼60% to ∼30% of the field of view. This trend is then reversed for the remainder of mitosis where net F-actin assembly outpaced net F-actin disassembly, leading to an increase in wave occupancy to ∼80% of the cortex before plateauing again at ∼60% of the field of view (Figure 3B’’-B’’’’).

### Cdk1 and APC/C activation are associated with changes to excitable dynamics

The above results suggest that the cell cycle regulates cortical excitability. To test this, we first incubated cells in the protein synthesis inhibitor, cycloheximide, to induce a late interphase arrest (Newport and Kirschner, 1984) due to suppression of cyclin synthesis and Cdk1 activation (Murray and Kirschner, 1989). Consistent with a role for cell cycle regulation of wave dynamics, cells incubated in 100 μg/mL cycloheximide develop robust (high amplitude, clearly spaced) cortical waves uninterrupted by the transitions associated with entry and exit from mitosis (Figure 4 A-B; Video 8).

**Figure 4.**
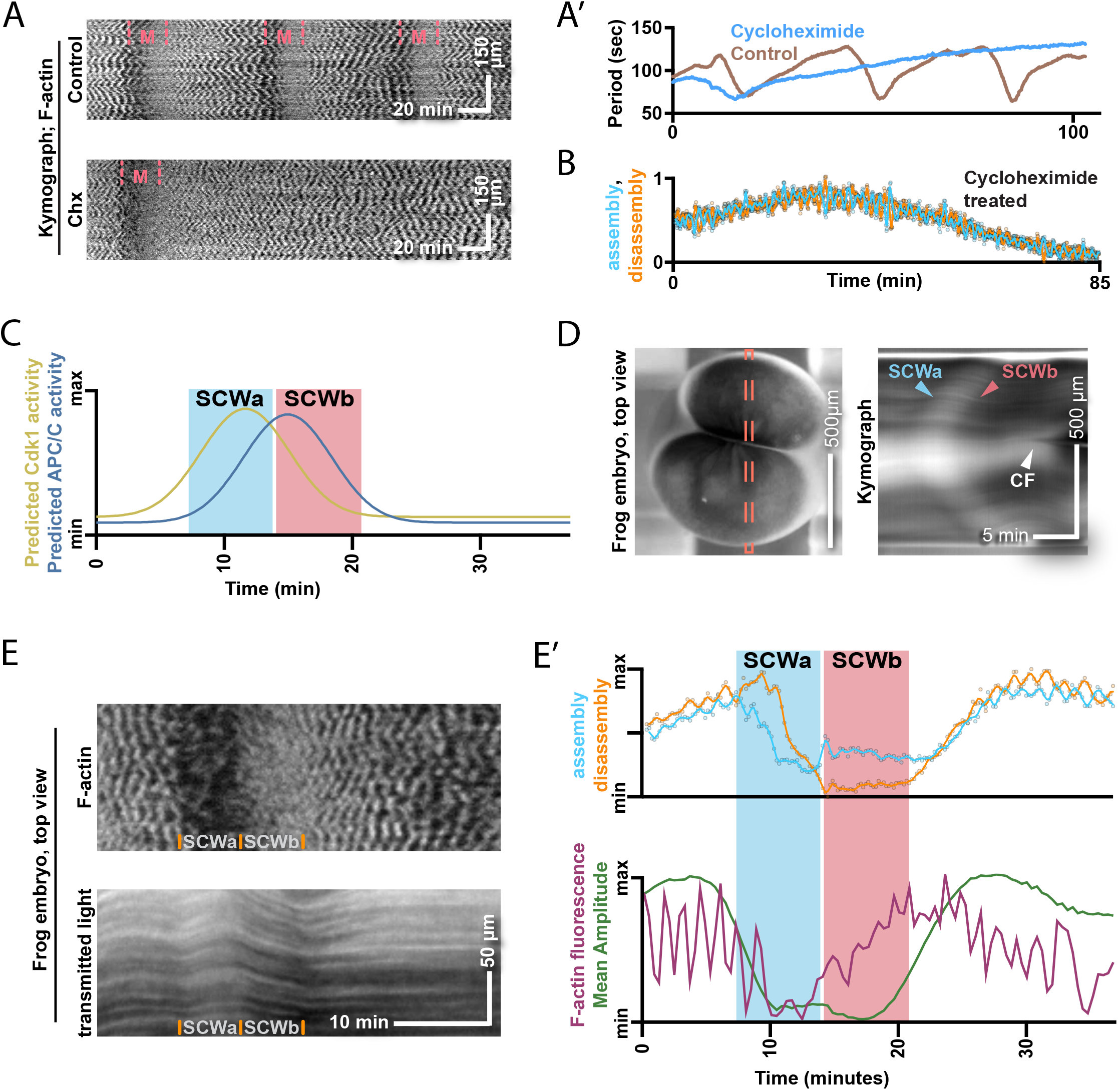
Changes in cortical excitability correlate with the appearance of the surface contraction waves. (A) Kymographs from a control embryo (top) and a cycloheximide-treated embryo (Chx, bottom). Both embryos were injected with Alexa647 conjugated UtrCH protein and treated with 3μM nocodazole to prevent cytokinesis. Entry into and out of mitosis are indicated with pink dashed lines; the cycloheximide-treated embryo goes through one round of mitosis immediately following cycloheximide treatment and then arrests. (A’) Plot showing the wave period over time from the embryos shown in A. The control embryo shows continuous fluctuations in wave period corresponding to cell cycle progression, while the cycloheximide-treated embryo asymptotically approaches a maximum wave period. (B) Plot showing total net F-actin assembly (light blue) and total net F-actin disassembly (light orange) over a 25 second window in a cyclohexim-ide-treated embryo. Points on the plot represent single measurements, solid lines represent a rolling average of 5 frames. At no point do the assembly and disassembly measurements significantly diverge from each other. (C) Schematic diagram illustrating the expected peaks in Cdk1 and APC/C activity relative to the two surface contraction waves (Rankin and Kirschner, 1997; Pérez-Mongiovi et al., 1998; Bischof et al., 2017). (D) Still image (left) and kymograph (right) of a cleaving frog embryo illustrating the changes in pigment distribution as the two surface contraction waves propagate across the surface of the embryos. The kymograph was drawn from the region indicated in the pink dashed box on the first still frame. The start points of the SCWs are marked with “SCWa” and “SCWb” and the appearance of the cleavage furrow following SCWb is marked with “CF”. (E) Kymographs from a nocodazole-treated embryo injected with Alexa647 conjugated UtrCH protein, the UtrCH fluorescence is shown on top, and transmitted 647 laser light (measuring pigment density) is shown on the bottom. The start and end points of SCWa and SCWb, as approximated from changes in pigment density, are indicated at the bottom of each kymograph. (E’) Plots superimposed over the estimated time frames for the start and end points of SCWa and SCWb, as approximated from changes in pigment density in E. The top plot shows the total net F-actin assembly (light blue) and total net F-actin disassembly (light orange) over a 36 second window in the embryo shown in E. Points on the plot represent single measurements, solid lines represent a second rolling average of 3 points. The amounts of total net assembly and disassembly diverge at the beginning of SCWa and reverse at the beginning of SCWb. The bottom plot shows a representative trace of UtrCH fluorescence in a ∼9μm2 ROI and the mean wave amplitude for the cell shown in E.

The finding that wave properties begin to reset approximately halfway through mitosis suggests that cortical excitability may be responsive to different stages of mitosis. Unfortunately, while cortical microtubules are a robust marker for mitotic entry and exit, they offer no information on the sub-stages of mitosis. To better understand how wave behavior is influenced by cell cycle state, we instead compared wave metrics to the surface contraction waves (SCWs). The SCWs are sequential waves of cortical relaxation (SCWa) and contraction (SCWb), which precede cytokinesis in large cells (Hara, 1971; Hamaguchi and Hiramoto, 1978; Hara *et al*., 1980; Amiel and Houliston, 2009). SCWa appears concurrent with Cdk1 activation, while SCWb appears concurrent with anaphase promoting complex/cyclosome (APC/C) activation (Rankin and Kirschner, 1997; Pérez-Mongiovi *et al*., 1998; Bischof *et al*., 2017; Figure 4C); thus, SCWs provide finer temporal markers for mitotic progression than cortical microtubules. In *X. laevis* embryos, the SCWs can be detected by virtue of the pigmentation at the animal hemisphere (Hara *et al*., 1980); during SCWa, the cortex relaxes, decreasing cortical pigment density and lightening the cell, while during SCWb the cortex contracts, increasing cortical pigment density and darkening the cell (Figure 4D; Video 9). For this analysis we used *X. laevis* zygotes injected with dye-labeled UtrCH protein and treated with nocodazole as embryos showed more robust surface contraction waves than activated oocytes, and nocodazole treatment ensured that a field of view would not be interrupted by a cleavage furrow but otherwise caused no obvious changes to wave dynamics. Consistent with our findings using cortical microtubules as a readout for cell cycle state, we found a tight correlation between F-actin wave dynamics and cell cycle state as inferred from SCW onset. Moreover, we found that the transition between SCWa (high Cdk1 activity) and SCWb (high APC/C activity) perfectly marked the transition between sparse F-actin waves and tightly packed F-actin waves (Figure 4E, E’).

### Changes in wave behavior scale with development and cell size

The early embryonic cleavages in *X. laevis* embryos continually change the cytoplasmic volume, the ratio of cell cortex to cytoplasmic volume, the ratio of DNA to cytoplasmic volume, and the overall complement of proteins, each of which have been shown to influence intracellular scaling during early development (Good *et al*., 2013; Hazel *et al*., 2013; Brownlee and Heald, 2019; Hara and Merten, 2015; Chen *et al*., 2019). Excitable dynamics are insensitive to cell size in some cases (Gerhardt *et al*., 2014; Xiao *et al*., 2017) and scale with cell size in other cases (Xiao *et al*., 2017). To determine whether features of cortical excitability scale with size in *X. laevis* embryos, we analyzed wave dynamics at different stages of early embryonic development. First, we used cycloheximide to arrest cells in interphase at different stages of development to standardize wave behavior between cells and measured two-dimensional wave area in cycloheximide arrested cells by thresholding UtrCH fluorescence (see methods). Wave area measurements were highly variable within and between cells using single time point analysis, but by averaging 100 sequential time points for each cell, we found a good agreement between cells at a given time point and a steady decrease in wave area from ∼30μm^2^ to ∼15μm^2^ as cells became smaller (Figure 5A). This finding agreed with our qualitative interpretation of individual still frames at different stages of development (Figure 5A’), suggesting that indeed excitable dynamics change as development progresses. Next, we asked whether the wave period showed a similar trend. By analyzing the wave period throughout multiple full cell cycles at different stages of development, we found a consistent increase and then decrease in both the maximum and minimum wave period throughout development, with the longest wave periods identified around stage 4-5 (16-32 cells; cell diameter 400-500μm; Supplemental Figure 5). As described above (Figure 3B), cells at all stages of development invariably showed some change in wave period throughout the cell cycle (Figure 3A’, 5B). However, the changes observed in smaller cells were consistently less extreme than those identified early in development. Indeed, by stage 6-6.5 (cell diameter ∼325 μm), most cells showed only subtle changes in either wave period or amplitude over the course of the cell cycle (Figure 5B; Video 10). To quantify these changes, we measured the difference between the maximum and minimum wave period and plotted this as a function of cell size (Figure 5C, 5D, Supplemental Figure 5). Similar to the wave area, measurements were irregular due to both biological variability and sample quality, but the overarching trend was a decrease in the variability of the wave period as development progresses. Finally, we asked whether cortical excitability was a consistent feature throughout embryogenesis. We easily identified F-actin waves in embryos as late as stage 9, but later stage embryos (∼stage 20) showed no evidence of excitable waves (Supplemental Figure 6A, A’).

**Figure 5.**
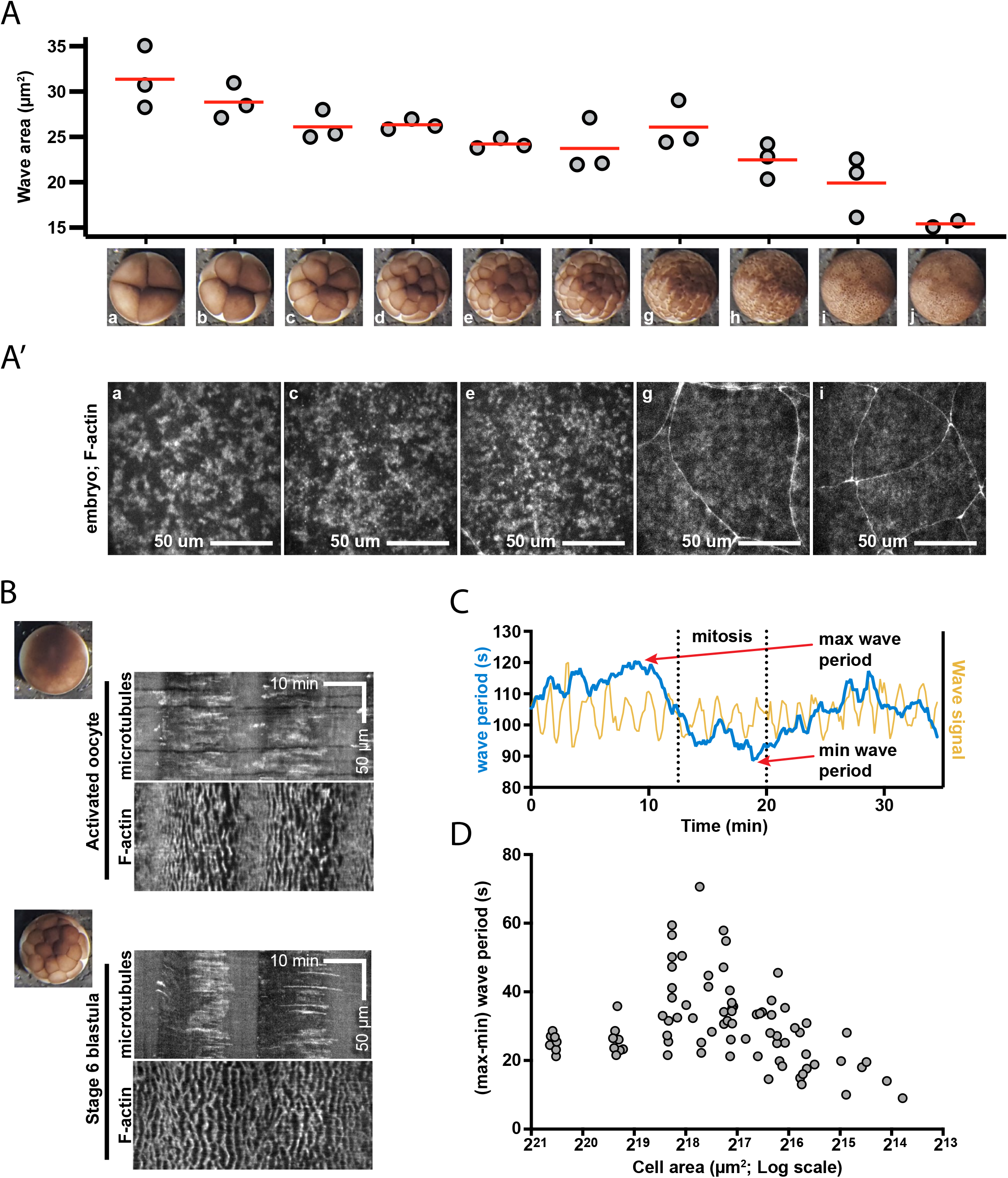
Cortical excitability scales with development and cell size. (A) Plot showing the two-dimensional area of individual waves throughout early development, as assessed by automated thresholding of the UtrCH signal. An example of each developmental stage assessed is shown below each column. The median wave area was measured in three embryos for each stage, the mean of these three embryos is indicated in red. (A’) Representative examples from five different developmental stages shown in A. (B) Kymographs from an activated oocyte (top) and stage 6 embryo (bottom) expressing mCh-EMTB (microtubules, top kymograph for each cell) and GFP-UtrCH (F-actin, bottom kymograph for each cell). An example of the developmental stages used are shown to the left of each kymograph pair. (C) Representative trace of UtrCH fluorescence (yellow) from a small ROI as a cell going through mitosis. The wave period is shown in blue, with the maximum and minimum wave period marked with red arrows. (D) Plot of the difference between maximum and minimum wave period throughout the cell cycle plotted against cell size as estimated from the simple multiplication of its longest and shortest axis. The x-axis is shown as a log scale to account for large changes in cell area early in embryogenesis.

### Excitable dynamics is detected within the cleavage furrow

F-actin waves were previously noted within the cleavage furrow of blastula-stage frog embryos (Bement *et al*., 2015). However, cortical excitability has never been investigated within the cleavage furrow of early-stage frog (stage 4-6) embryos. Moreover, the relationship between cleavage furrow cortical excitability and non-furrow excitability is unclear. Thus, we sought to detect and characterize wave parameters within the cleavage furrow of early-stage *X. laevis* embryos. Consistent with previous work, we detected traveling waves of F-actin assembly and disassembly within the cleavage furrow of early (stage 5) frog embryos. However, these waves were low in amplitude relative to the excitable F-actin waves outside of the furrow and were largely obscured by the thick cables of F-actin that make up the contractile ring (yellow arrowheads, Figure 6A; Video 11, Video 12). Because the cables were stable over space and time relative to the traveling waves, calculating the change in fluorescence over time in each pixel (also known as a difference subtraction; Bement *et al*., 2015; Landino *et al*., 2021) allowed better visualization and quantification of waves in the cleavage furrow (Figure 6A, A’; Video 11, Video 12). When compared to non-furrow waves, there was no significant difference in the wave period or temporal width (Figure 6B, C). However, the rate of new F-actin assembly (i.e., the wave amplitude following a difference subtraction) of furrow waves was significantly lower than the rate of assembly of non-furrow waves (Figure 6D). Thus, while some wave properties are preserved between the furrow and non-furrow regions, the two sets of waves are spatially segregated and display both qualitatively and quantitively different properties.

**Figure 6.**
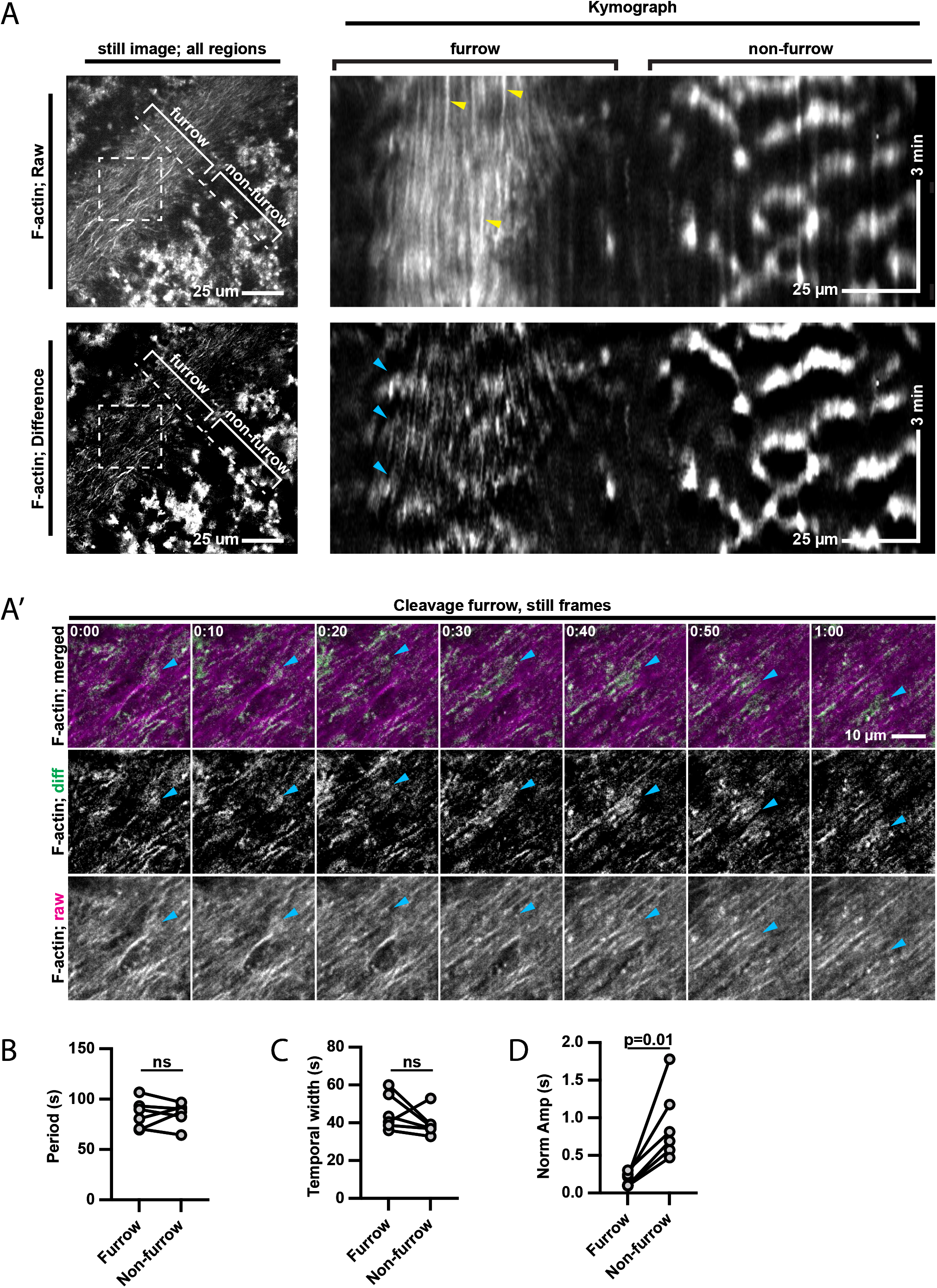
Excitable dynamics are detected within the contractile ring. (A) Still frames (left) and kymographs (right) from a raw (top) or a 30-second difference processed (bottom) time lapse data set. The kymographs are drawn from the region indicated with a dashed line on the respective still frames. Furrow and non-furrow regions are indicated on both still frames and kymographs. Yellow arrowheads indicate stable F-actin cables in the raw kymograph, blue arrowheads indicate waves in the difference kymograph (A’) Image montage derived from the boxed region marked with the dashed box in A. The raw images are shown on the bottom panel in grayscale and the top panel in magenta. The 30-second differenced images are shown in the middle panel in grayscale and the top panel in green. Blue arrowheads follow a propagating wavefront. (B-D) Plot of the wave period (B), temporal width (C), or normalized wave amplitude (D) from within the furrow region or non-furrow region. Each pair of points represents the mean furrow and non-furrow measurements from a single cell. Statistical significance for each measurement is marked as either not statistically significant (ns) or p=0.01 using a paired t test.

## Discussion

Cortical excitability is a consistent feature of embryos (Bement *et al*., 2015; Maître *et al*., 2015; Michaux *et al*., 2018), but the developmental genesis of cortical excitability is not well understood, nor is the relationship of cortical excitability to cell cycle progression, at least in *Xenopus*. To address these gaps, we explored and carefully quantified the origin and behavior of cortical excitability in activated *X. laevis* oocytes and embryos. This analysis revealed previously unnoticed cortical behaviors over both short time scales (minutes), as well as over developmental time (hours), and underlined the differences between different populations of excitable waves, namely those within the cleavage furrow and those outside of it.

Previous research in starfish oocytes revealed clear traveling waves prior to meiosis-I and meiosis-II (Bement *et al*., 2015), suggesting that this behavior was related to normal polar body emission. While we detected wave-like flickers of F-actin and a large increase in the overall amount of F-actin turnover during meiosis, these dynamics were not consistent or robust enough in *X. laevis* oocytes to confidently label them as excitable. Rather, the first clear examples of cortical excitability – unambiguous and uniform F-actin waves – were detected throughout the cell cortex about 15 minutes after egg activation. Previous work on cortical excitability in *X. laevis* embryos hinted at potential changes in wave dynamics that coincided with cell cycle progression (Bement *et al*., 2015). Here, using improved and complementary imaging approaches, we show that quantitative wave metrics are in a continuous state of change and reset with each passage through the cell cycle. Specifically, the wave period steadily increased throughout interphase, reaching peak values (∼100 seconds) at the beginning of the next mitosis. Wave amplitude and the fraction of the cortex occupied by waves both peaked early in interphase and either remained steady or declined slightly until the next mitosis. The most dramatic changes to wave behavior coincided with appearance of the relaxation (SCWa) and contraction (SCWb) waves preceding cytokinesis. That is, the fraction of the cortex occupied by waves fell by approximately half concurrent with the appearance of the relaxation wave (SCWa), leading to sparse F-actin wave distribution throughout the cortex. This behavior reversed coincident with the contraction wave (SCWb), where F-actin waves rapidly expanded into a dense network off waves. Biochemically, the appearance of SCWa is linked to Cdk1 activation while SCWb is linked to APC/C activation (Rankin and Kirschner, 1997; Pérez-Mongiovi *et al*., 1998; Bischof *et al*., 2017). Finally, treatment with cycloheximide arrests cells in interphase with low Cdk1 activity; this treatment prevented fluctuations in wave behavior over time, further establishing a causal link between wave behavior and cell cycle state.

The interplay between cortical excitability and the cell cycle presented above provides an interesting contrast to those from earlier studies. In mast cells, randomly oriented cortical waves were converted to bullseye and spiral waves during mitosis in some cells, and completely lost in others (Xiao *et al*., 2017). In starfish, excitability was consistently and completely lost during early M-phase and reappeared shortly after anaphase onset (Bement *et al*., 2015). In contrast, in the frog system, where cortical excitability manifests continuously, it nonetheless reproducibly fluctuates according to cell cycle state. This observation underscores a point suggested previously (Michaud *et al*., 2021), namely, that the maximal “all-or-none” response – that is, the resulting wave – of cortical excitability can encompass a spectrum of different behaviors depending on the cellular environment. This lies in contrast to more extreme examples of excitability, such as neuronal action potentials, where the maximal response is typically consistent within the same cell. While both F-actin waves and the electrochemical waves of neurons are governed by the same principles, from the biological standpoint, this difference makes sense. Neurons rapidly send information in the form of undamped waves with the eventual interpretation of the information reflecting the participation of many cells. Cortical excitability, in contrast, must be interpreted within a single cell and therefore the interpretation – that is, the cellular response – to cortical excitability must be fine-tuned in space and time according to the needs of the cell.

During early development, the effect of cell cycle state on cortical excitability was so pronounced that entry into and out of mitosis could be deduced by eye in the absence of any other markers. However, as embryos continued to develop, we found that wave behavior became less and less responsive to cell cycle state. Eventually, entry into and out of mitosis could only be detected with a dedicated cell cycle marker or by careful quantification of wave dynamics. Simultaneously, we found that two-dimensional wave area scaled with decreasing cell size as development progressed from stage 2 (two cells) to about stage 8 (∼4,000 cells). We eventually lost the ability to detect excitability at all, suggesting that cortical excitability is primarily functional during early embryonic development. However, it is possible that additional changes in wave behavior occur between the period of time where we focused our attention (∼stage 2 to ∼stage 8) and the developmental period where we found no evidence for cortical excitability (∼stage 20). The observation that cortical excitability scales with developmental progression is intriguing both from a mechanistic as well as a functional perspective. While investigating the mechanism of scaling is beyond the scope of the present work, we find it likely that previously proposed mechanisms for cellular scaling, such as changes to the surface to volume ratio or nuclear to cytoplasmic ratio, could influence the amounts or activity of the relevant players at the cell cortex (for review see Miller *et al*. 2020).

While it remains difficult to assign a specific biological function to cortical excitability in *X. laevis* embryos, we are intrigued that it predominantly manifests in large cells, which face biophysical challenges that small cells do not. For example, *X. laevis* embryos are far too large for diffusion to be an effective means of signaling. Instead, much of the early coordination for cell division is driven by trigger waves of Cdk1 activation which coordinate mitotic entry across the ∼1 mm diameter cell (Chang and Ferrell, 2013). One role of this coordinated trigger wave is to regulate the dynamics of microtubules, which ultimately delivers cleavage signals to the cell cortex (Ishihara *et al*. 2014). Similarly, it may be the case that excitability of the cell cortex allows it to respond to Cdk1 activation and inactivation and remodel the vast cortical cytoskeleton in a rapid and coordinated fashion. Indeed, we found that during interphase, the rate of total net F-actin assembly and disassembly were largely matched. However, these trends rapidly diverged upon Cdk1 activation – as determined by SCWa onset – with net assembly dropping and total net disassembly rising. This divergence led to the net disassembly of ∼50% of the total wave area over the course of ∼60 seconds (see above). This trend immediately reversed upon APC/C activation – as determined by SCWb onset – with total net assembly outpacing total net disassembly and cortical F-actin transitioning to a dense network of waves. The finding that changes in net F-actin assembly/disassembly rates closely match the appearance of the surface contraction waves suggest that these rapid remodeling events may be necessary for the SCWs to form properly. Consistent with this line of thought, we found that wave dynamics became less sensitive to cell cycle state over time, mirroring the decreasing intensity of the surface contraction waves. While it is conceivable that a linear system could accomplish the same feats, an excitable cell cortex makes it uniquely posed to execute these steps faster for two simple reasons. First, the intertwined positive and negative feedback loops of excitable systems quickly integrate and amplify small stimuli. Second, as shown here, excitable waves are in a continuous state of remodeling, with each wave completely turning over in approximately one-half wave period (30-60 seconds), which allows for rapid replacement of existing structures. Thus, an excitable cell cortex may be an evolutionary adaptation allowing the comparatively vast cell cortex of early blastomeres to rapidly remodel according to the changing demands of early embryogenesis.

In addition to the remodeling events observed during the SCWS, we also observed a rapid transition from baseline cortical excitability to the formation of the cleavage furrow during cytokinesis. Strikingly, this event resulted in the formation of a new population of F-actin waves confined to the cleavage furrow region that are clearly segregated in space from the otherwise pan-cortical actin waves. While the furrow waves were similar to non-furrow waves in terms of period and temporal width, the F-actin waves within the cleavage furrow were significantly lower in amplitude than the non-furrow waves. Moreover, the furrow waves appeared to be propagating within a background of relatively stable F-actin cables. Altogether, the furrow waves appeared to represent a quantitatively different regime of cortical excitability.

Our previous analysis of cortical excitability in *X. laevis* revealed low amplitude waves of active Rho associated with the pan-cortical F-actin waves. This work suggested that the Rho GEF Ect2 was a critical component for regulating the amplitude and behavior of the Rho waves, and that Ect2 accumulation at the equatorial cortex during cytokinesis is responsible for mediating the transition between a state of excitable Rho dynamics and a state with largely stable Rho activity in the furrow (Bement *et al*., 2015, Goryachev *et al*., 2016). Functionally, this model suggests that cortical excitability could represent an “idling” of the cell cortex, allowing it to quickly respond to faint cues from the mitotic spindle. As previously proposed, it is possible that the mutually exclusive wave patterns (furrow versus non-furrow) we observed could be a consequence of locally concentrating the Rho GEF Ect2 at the cell equator and depleting it from the neighboring areas (Bement et al. 2015). Alternatively, it is possible that this mutually exclusive pattern is a consequence of different wave regulators – e.g., different GTPases, F-actin assembly factors, or F-actin binding proteins – that could be active in the waves outside of the contractile ring and excluded from within the contractile ring or vice versa. Consistent with this possibility, Arp2/3 networks have been proposed to antagonize Rho signaling during cytokinesis (Pal *et al*., 2020), and inactivation of Rac at the cleavage plane by the centralspindlin complex is necessary for normal cytokinesis (Canman *et al*., 2008; Bastos *et al*, 2012; Zhuravlev *et al*., 2017), making the segregation of these players a prerequisite for a succsful cytokinesis. GTPases are ideal players in excitable systems because they can engage in both positive feedback and delayed negative feedback. Accordingly, we are intrigued by the possibility that segregation of different GTPases could explain the different behavior of the furrow and non-furrow waves. While the possibilities discussed above are not mutually exclusive, untangling the origin of the differences between the cleavage furrow and non-furrow waves is essential to understanding how cortical excitability contributes to cytokinesis.

## Video Legends

**Video 1**. Corresponds to Figure 1A. *Xenopus laevis* embryo expressing GFP-UtrCH. Video illustrates robust cortical waves of F-actin (cortical excitability) in a stage 5 *X. laevis* embryo. Time is in min:sec.

**Video 2**. Corresponds to Figure 1B. *Xenopus laevis* oocyte expressing GFP-UtrCH (F-actin; left) and mCh-EMTB (microtubules; right). Video shows changing F-actin dynamics as the oocyte passes through meiosis-I (minutes 196 to 307) and arrests in meiosis-II (minutes 341 and later). Time is in min:sec.

**Video 3**. Corresponds to Figure 2A. Video shows a mature (meiosis-II arrested; left) and a recently activated (enter interphase; right) *Xenopus laevis* oocyte. Both oocytes are expressing GFP-UtrCH. Time is in min:sec.

**Video 4**. Corresponds to Figure 2B. *Xenopus laevis* oocyte expressing GFP-UtrCH. Video shows activation response as the mature oocyte is exposed to the calcium ionophore ionomycin. The oocyte is mature (meiosis-II arrested) at the beginning of the video. The time stamp counts down to 00:00 when the cell begins to respond to the ionomycin flowed into the imaging chamber. Ionomycin is flowed in at 01:10 (counting down) and flushed out at 3:55 (counting up). Time is in min:sec.

**Video 5**. Corresponds to Figure 3A. *Xenopus laevis* embryo injected with Alexa^647^-conjugated UtrCH protein. Video illustrates the changing cortical F-actin dynamics preceding each round of cytokinesis. Time is in min:sec.

**Video 6**. Corresponds to Figure 3B. Activated *Xenopus laevis* oocyte expressing GFP-UtrCH (F-actin; left) and mCh-EMTB (microtubules; right). Videos illustrates the changing cortical F-actin dynamics as the cell passes through mitosis. Mitosis, as judged by the loss of cortical microtubules, starts at 10:30. Mitotic exit, as judged by the repopulation of cortical microtubules, begins at 18:15. Time is in min:sec.

**Video 7**. Corresponds to Figure 3B’’. Video shows spatial map of GFP-UtrCH loss and gain on the left, and mCH-EMTB on the right. Regions where net F-actin has increased in the previous 60 seconds are shown in blue, regions where net F-actin has decreased in the previous 60 seconds are shown in orange. Mitosis, as judged by the loss of cortical microtubules, starts at 9:05. Note that following this time point there is significantly more area reporting a loss of GFP-UtrCH signal than area reporting a gain in GFP-UtrCH signal (corresponds to the second pink arrowhead in Figure 3B’’). Mitotic exit, as judged by the repopulation of cortical microtubules, begins at 17:20. Time is in min:sec.

**Video 8**. Corresponds to Figure 4A. Video shows two nocodazole-treated *Xenopus laevis* embryos injected with Alexa^647^-conjugated UtrCH protein. The embryo on the left was otherwise untreated, and the embryo on the right was treated with 100μg/ml cycloheximide. The embryo on the left goes through three rounds of mitosis starting at time stamps 06:00, 41:20 and 75:00, and appears to be entering a fourth round starting at time stamp 106:00. The cycloheximide-treated cell goes through one round of mitosis starting at time stamp 06:00, after which it arrests in interphase. Time is in min:sec.

**Video 9**. Corresponds to Figure 4D. Video shows a *Xenopus laevis* embryo passing through its first mitosis under a dissecting microscope. The kymographs unfolding as video plays illustrates the appearance of the two surface contraction waves and the cleavage furrow. Time is in min:sec.

**Video 10**. Corresponds to Figure 5B. Video shows a stage 6 *Xenopus laevis* embryo (left) and an activated oocyte (right), both expressing GFP-UtrCH (F-actin; top), and mCh-EMTB (microtubules; bottom). Video illustrates the differences in mitotic F-actin dynamics between cells early in development and later in development. Time is in min:sec.

**Video 11**. Corresponds to Figure 6A. Video shows a *Xenopus laevis* embryo injected with Alexa^647^-conjugated UtrCH protein. The raw data are shown on the left, a 30-second difference movie (subtracting out unchanging or decreasing signal from the previous 30 seconds) is shown on the right. Video illustrates the presence of F-actin waves within the contractile ring. Time is in min:sec.

**Video 12**. Corresponds to Figure 6A. Video shows a *Xenopus laevis* embryo injected with Alexa^647^-conjugated UtrCH protein. The raw data are shown in magenta, a 30-second difference movie (subtracting out unchanging or decreasing signal from the previous 30 seconds) is shown in green. Video illustrates the subtraction of large station cables, and shows a magnified view of the cleavage furrow relative to Video 11. Time is in min:sec.

## Materials and Methods

### Embryo preparation

Mature *Xenopus laevis* females were injected in the dorsal lymph sac with 800 units of human chorionic gonadotropin (HCG; MP Biomedicals) and incubated at 18°C for 12–18 h before use. Eggs were laid into 1X high-salt Modified Barth’s Saline (high-salt MBS; 108mM NaCl, 0.7mM CaCl_2_, 1mM KCl, 1mM MgSO_4_, 5mM HEPES, 2.5mM NaHCO_3_) and fertilized *in vitro* with testes fragments. 15-30 minutes following fertilization, embryos were dejellied in a solution of 2% cysteine in 0.1X Marc’s Modified Ringer’s (MMR; 100 mM NaCl, 2 mM KCl, 1 mM MgCl2, and 5 mM HEPES, pH 7.4) and then rinsed thoroughly with 0.1X MMR. Embryos were injected once at the 1-cell stage (30-45 minutes following fertilization) in 5% Ficoll (Sigma) in 0.1X MMR with 5nl of *in-vitro* synthesized mRNA or purified protein and maintained in 0.1X MMR during development.

### Oocyte preparation and activation

Approximately half ovaries were removed from mature *Xenopus laevis* females anaesthetized with MS-222. Ovary chunks were cut into ∼5mm pieces, rinsed thoroughly in 1X Barth’s solution (87.4 mM NaCl, 1 mM KCl, 2.4 mM NaHCO_3_, 0.82 mM MgSO_4_, 0.6 mM NaNO_3_, 0.7 mM CaCl_2_ and 10 mM HEPES at pH 7.4) and then treated with 8 mg/ml collagenase in 1X Barth’s solution for 1 hour at 17°C. After rinsing thoroughly with 1X Barth’s solution, oocytes were allowed to recover overnight at 17 °C. Following recovery, follicle cells were manually removed with forceps and oocytes were injected with 40nl of *in-vitro* synthesized mRNA or purified protein. Following injection, oocytes were treated with 5μg/ml progesterone in 1X Barth’s for 15 minutes, and then incubated in 1X Barth’s overnight at 17°C. The following day, mature eggs were activated by soaking in 10μg/ml ionomycin in 0.1X MMR for 30-60 seconds. Activated eggs were maintained in 0.1X MMR while imaging. For the oocytes in Figure 1B, oocytes were maintained at 17°C overnight following injection and then treated with progesterone shortly before imaging.

### Constructs, mRNA synthesis, and microinjection

Plasmids for eGFP-UtrCH, mCherry UtrCH, 3xGFP-EMTB, and mCherry-EMTB were made as described previously (Burkel *et al*. 2007; Miller and Bement 2009). Plasmids were linearized following the open reading frame and used as a template for mRNA synthesis using the mMessage Machine SP6 kit (Ambion), mRNAs were polyadenylated using a Poly(A) tailing kit (Ambion) and purified with a phenol-chloroform extraction and isopropanol precipitation. Alexa Fluor® 647-conjugated UtrCH protein was generated as described previously (Landino *et al*. 2021). eGFP-UtrCH and mCherry-UtrCH mRNA were injected into oocytes at a needle concentration of 20ng/nl, into embryos at 80ng/nl for imaging stages 4-6, and at 40ng/nl for imaging stages 6-8. mCherry-EMTB mRNA was injected into oocytes at a needle concentration of 50ng/nl. 3xGFP-EMTB mRNA was injected into embryos at a concentration of 500ng/nl for imaging stages 4-6.

### Drug treatments

Embryos were arrested in interphase by in incubating in 0.1X MMR with 100ug/ml cycloheximide. Embryos were treated with nocodazole by soaking in 1μg/ml (3.3μM) nocodazole.

### Microscopy and image processing

For all imaging experiments, embryos were maintained in 0.1X MMR at room temperature. For all confocal experiments, embryos or mature/activated oocytes were gently compressed with a clean #1.5 coverslip. Oocytes were activated on the microscope (Figure 2B) by mounting oocytes in a custom metal slide with a flow channel that allowed drugs to be flowed into and out of the chamber. The images in Figure 4D and the reference images in Figure 5A and 5B were acquired using a Samsung Galaxy S6 camera mounted to an Olympus SZX12 stereomicroscope with a Gosky Universal Cell Phone Adapter. The images in Figures 1, 2, 3B, 3B’’’’, 5, 6, and Supplemental Figures 1, 2 and 6 were collected using a CFI Plan Apo 60X/1.4 NA oil immersion objective on a Nikon Eclipse Ti inverted microscope equipped with an Opterra swept-field confocal unit (Bruker) using a 60μm pinhole array, a PZ-2000FT series piezo XYZ stage (ASI), and an Evolve Delta EMCCD camera (Photometrics). The microscope was controlled, and images acquired using Prairie View software (Bruker). The images in Figures 3A, 4A, 4E, and Supplemental Figure 5 were acquired using a UPlanSApo 10X/0.40 NA dry objective on an Olympus Fluoview 1000 laser scanning confocal. Z-series were acquiring using 0.5um steps (Bruker) or 5μm steps (Olympus), all confocal images shown are maximum intensity projections. Transmitted light images in Supplemental Figure 3 were acquired by measuring transmitted 640nm laser light. Still frames showed in Figures 1B, 2A, 2B, and 5B were from data registered for 2D drift using the StackReg plugin (Thévenaz *et al*., 1998) set to “translation” mode. Images in Figures 1B, 2A, 4A, 4E, and S6A were corrected for bleaching to optimize display by using an exponential fit within FIJI. Temporal color coding for Figure 1A’ was performed manually. Temporal color coding for Figure 1B’’ was performed using the temporal color code plugin packaged with FIJI (Kota Miura; Centre for Molecular and Cellular Imaging, EMBL Heidelberg, Germany). The kymographs in Figures 2A, 3A, 3B, 6A, and S6A’ were scaled in the y-axis using bicubic interpolation. All kymographs represent the average pixel intensity within a ∼2-16μm-thick slab.

### *Image quantification* and statistical tests

The workflow for wave amplitude, period, and width calculations were conceptually based off of a MATLAB framework written by Marcin Leda and Andrew Goryachev (Bement *et al*., 2015), and was reimagined here by Zac Swider and Ani Michaud in Python form to increase speed, accuracy, and access. Briefly, datasets were analyzed by breaking a field of view into n ∼5.5μm^2^ squares and using the mean signal within each box to quantify the wave amplitude, width, and period within each box (Supplemental Figure 1). Boxes were then averaged to approximate the population mean (Supplemental Figure 1B). Rolling analysis of long movies was accomplished by independently analyzing sub-sections of a movie. For example, the data in Figure 3B were generated by analyzing a 1,500-frame dataset in 1,451 50-frame sub-sections, each of which progressed by 1 additional frame (Supplemental Figure 1C).

Wave periods (Supplemental Figure 1A) were estimated by generating a full autocorrelation of each box mean using *numpy*.*correlate* and identifying the position of the first non-zero peak using *scipy*.*signal*.*find_peaks*. Wave maximum and minimum values were calculated by first smoothing the mean signal within a box with a Savitzky-Golay filter using *scipy*.*signal*.*savgol_filter*, and then identifying peak indices using *scipy*.*signal*.*find_peaks*. Temporal width of peaks was calculated using *scipy*.*signal*.*peak_widths* at full width half max. As multiple peaks were typically identified within each time point, the average of each peak metric was reported for each box.

The relative rates of net F-actin assembly and disassembly (Figures 3B’’ and Figure 5B, E’) were quantified using a custom script written in Python 3. Briefly, every frame was compared to the frame n time points previous to it. If the change in signal was positive for a pixel, it was considered to reflect net F-actin assembly. If the change in signal was negative for a pixel, it was considered to reflect net F-actin disassembly. The total amount of net F-actin assembly and disassembly for each time was calculated by averaging the total amount of positive and negative boxes for each time point. The percent occupancy of F-actin waves for a given field of view was calculated by thresholding the F-actin signal in FIJI and measuring the total area above the threshold. Two-dimensional wave area was approximated by thresholding the F-actin signal for every time point of a time series in FIJI and measuring composite ROIs, excluding particles on the edge of the image. The ROIs areas throughout the time series were averaged to cut down on noisy measurements. Difference movies were generated within FIJI by calculating the positive difference between n frames, typically (10-30 seconds). Figures 6B’, 6C, and 6D use unpaired Student’s t-test with α = 0.0S.

## Acknowledgements

We are grateful to our colleagues for continuous feedback on experiments, analysis, and data visualization. In particular we are grateful to Ann Miller for a detailed and thoughtful discussion of the manuscript. This work was supported by an award from the National Institutes of Health of Health (RO1GM052932) and from the National Science Foundation (MCB-1614190).

---

**Supplemental Figure 1.**
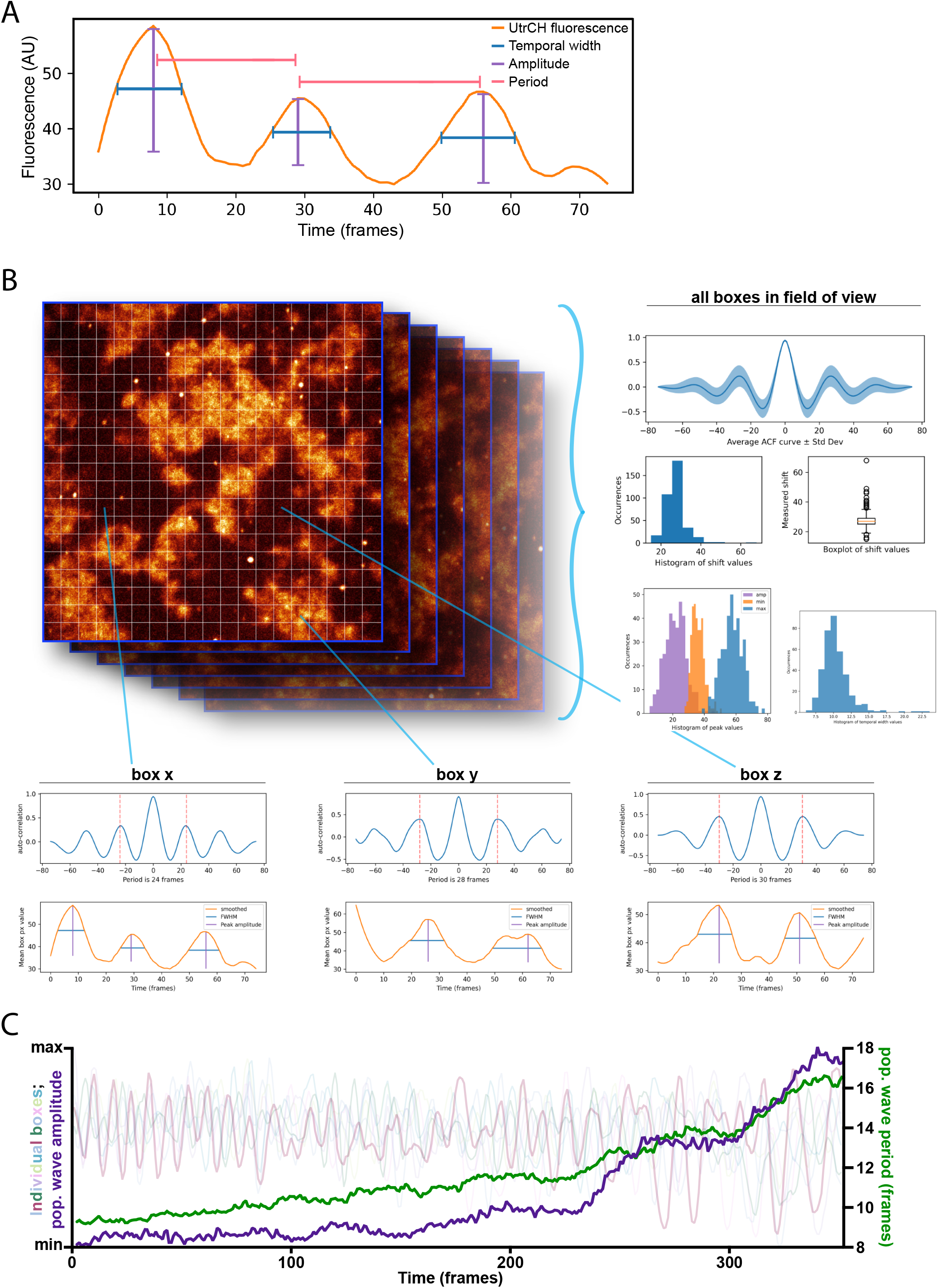
Processing and output from Python scripts. (A) Representative example of the UtrCH signal (orange) from a small region (5-8μm2) over time. The measurements for wave period, amplitude, and temporal width are shown in pink, purple, and blue respectively. (B) Schematic example of the workflow used to process a single short time-lapse. The image stack is broken in n squares with the same image depth as the parent stack, which are analyzed individually (below). In each box, the period is calculated using an autocorrelation. The wave amplitude and temporal width and are similarly calculated within each box by identifying signal peaks and calculating the peak height (peak-trough difference) and full width at half max of each peak. After each box is analyzed, the population of measurements (right) can be used to determine the average wave period, amplitude, and temporal width. (C) Example traces (in the background) of UtrCH fluorescence from some boxes in dataset where box the wave period and amplitude are increasing over time. Average measurements can be determined over time by sampling consecutive sub sections of the continuous datasets as illustrated in B.

**Supplemental Figure 2.**
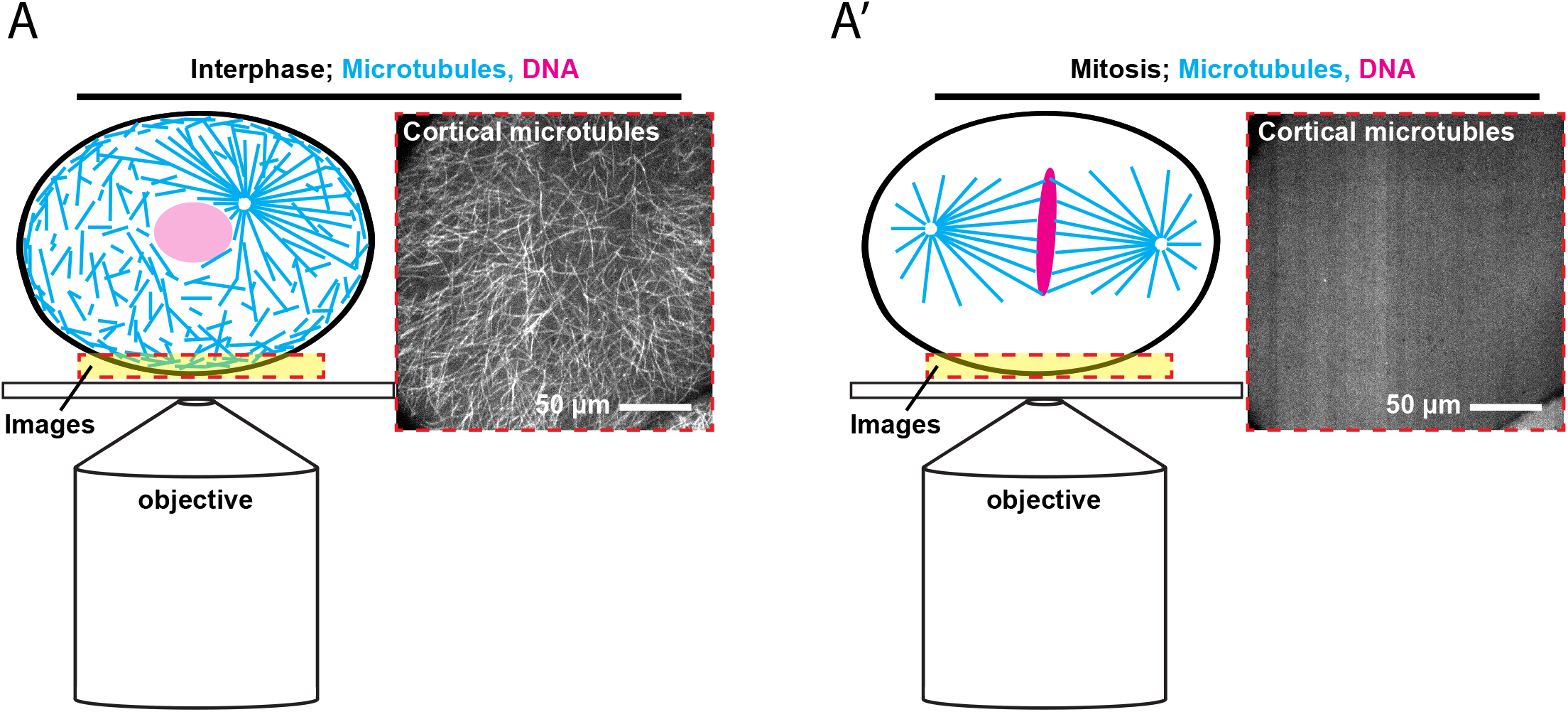
Behavior of cortical microtubules during interphase and mitosis. Schematic drawings of expected microtubule distribution in interphase (A) and mitotic (A’)cells, accompanied by representative z-stacks through the cell cortex (region showed boxed in red in schematic drawings) of the same cell during both interphase and mitosis.

**Supplemental Figure 3.**
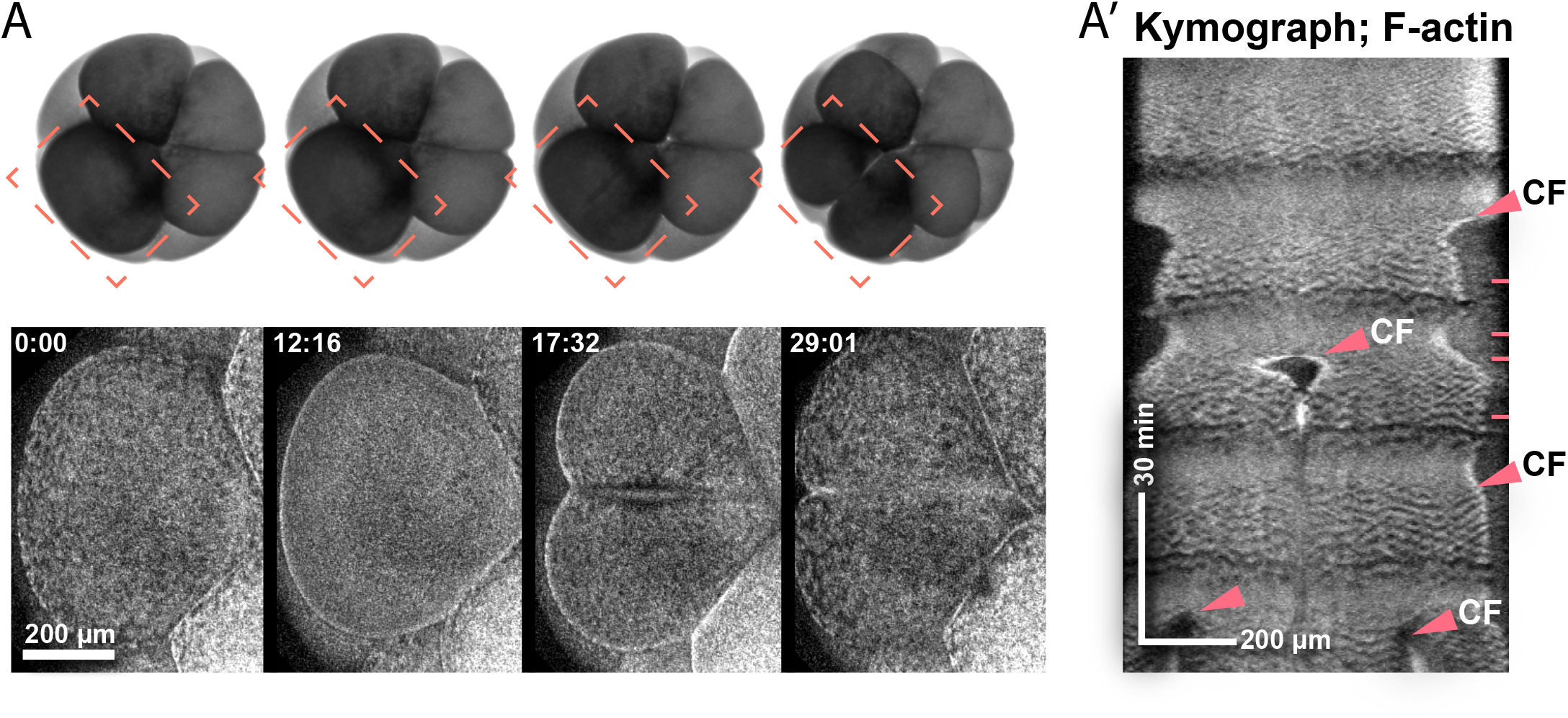
Low-magnification imaging of UtrCH during early embryogenesis. (A) Transmitted light (top) and UtrCH fluorescence (cropped, bottom) stills from a time lapse of an embryo cleaving from the 8-cell to the 16-cell stage showing changes in macro scale F-actin distribution during different cell cycle stages. (A’) Kymograph of the dataset shown in A. Appearance of cleavage furrows in the kymograph are indicated with arrowheads. Dashed lines indicate the time points used to generate the still images in A.

**Supplemental Figure 4.**
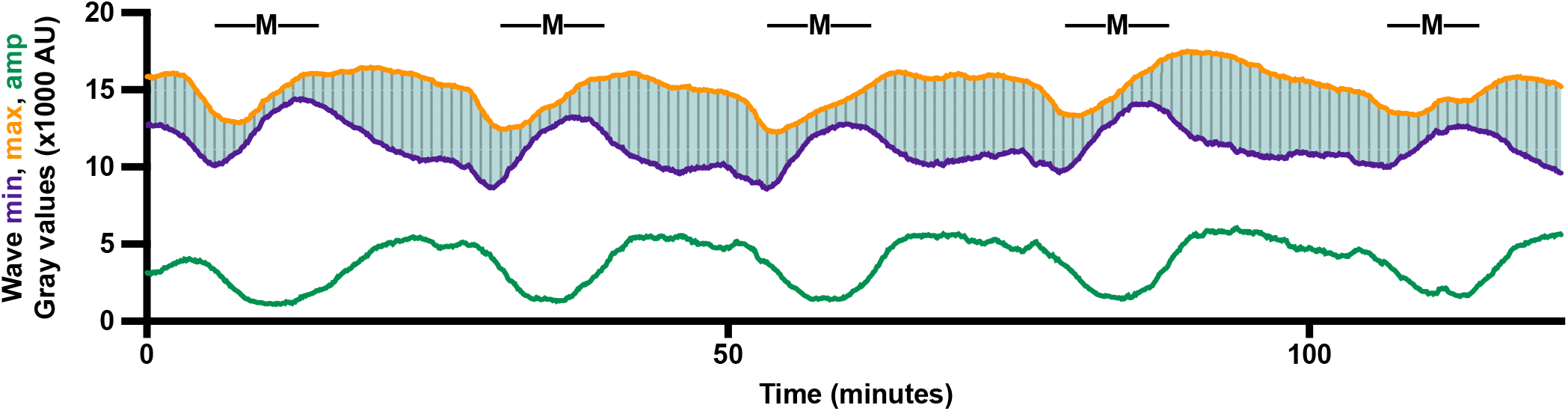
Relative changes in wave peak and trough values during mitosis. (A) Alternative visualization of the wave amplitude dataset shown in Figure 3B’. Wave max and min values are shown in orange and purple respectively. The wave amplitude is shown filled in between the two lines, and also shown below in green. Entry into and out of mitosis is shown with horizontal lines as in 3B’.

**Supplemental Figure 5.**
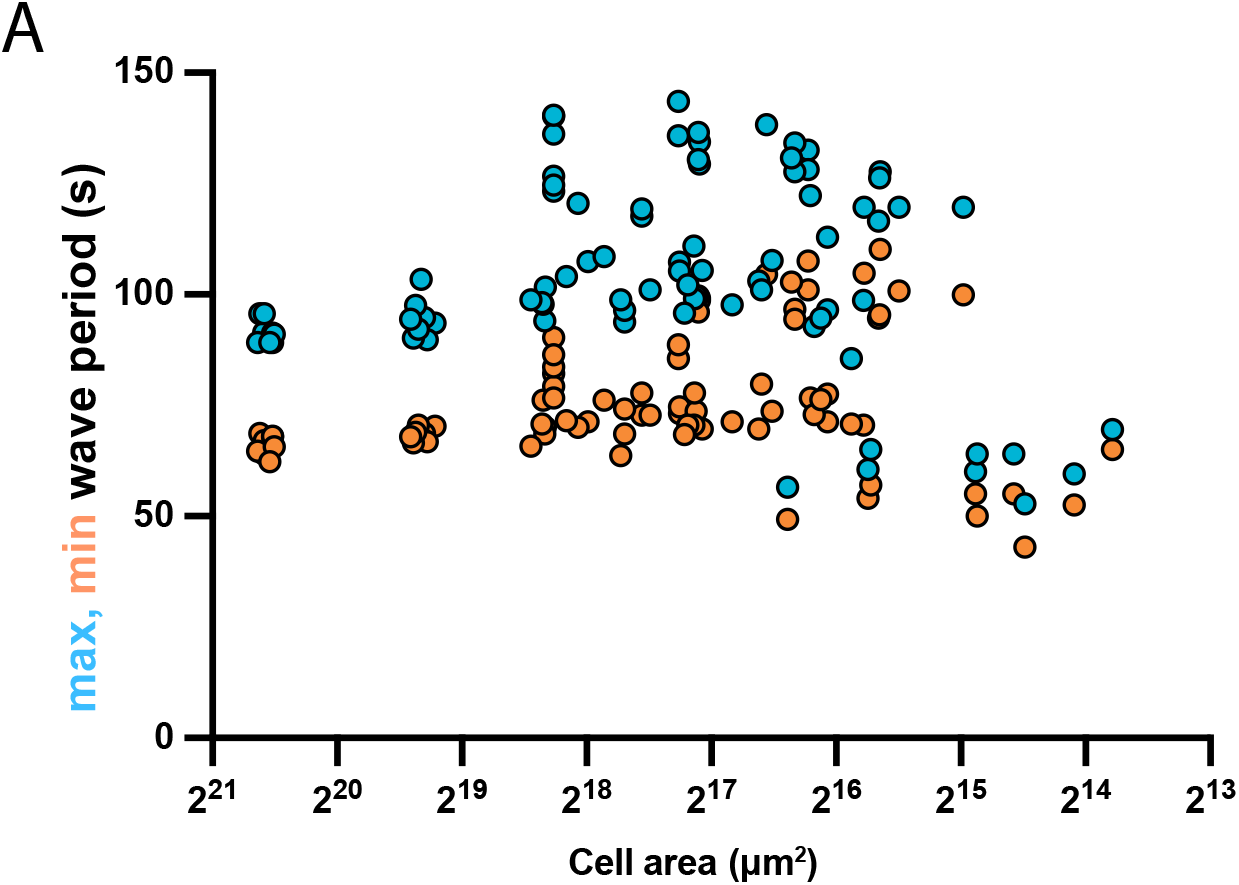
Wave period scales with cell size. Alternative visualization of the dataset shown in Figure 5D. Plot of the maximum and minimum wave period throughout the cell cycle plotted against cell size as estimated from the simple multiplication of its longest and shortest axis. The x-axis is shown as a log scale to account for large changes in cell area early in embryogenesis.

**Supplemental Figure 6.**
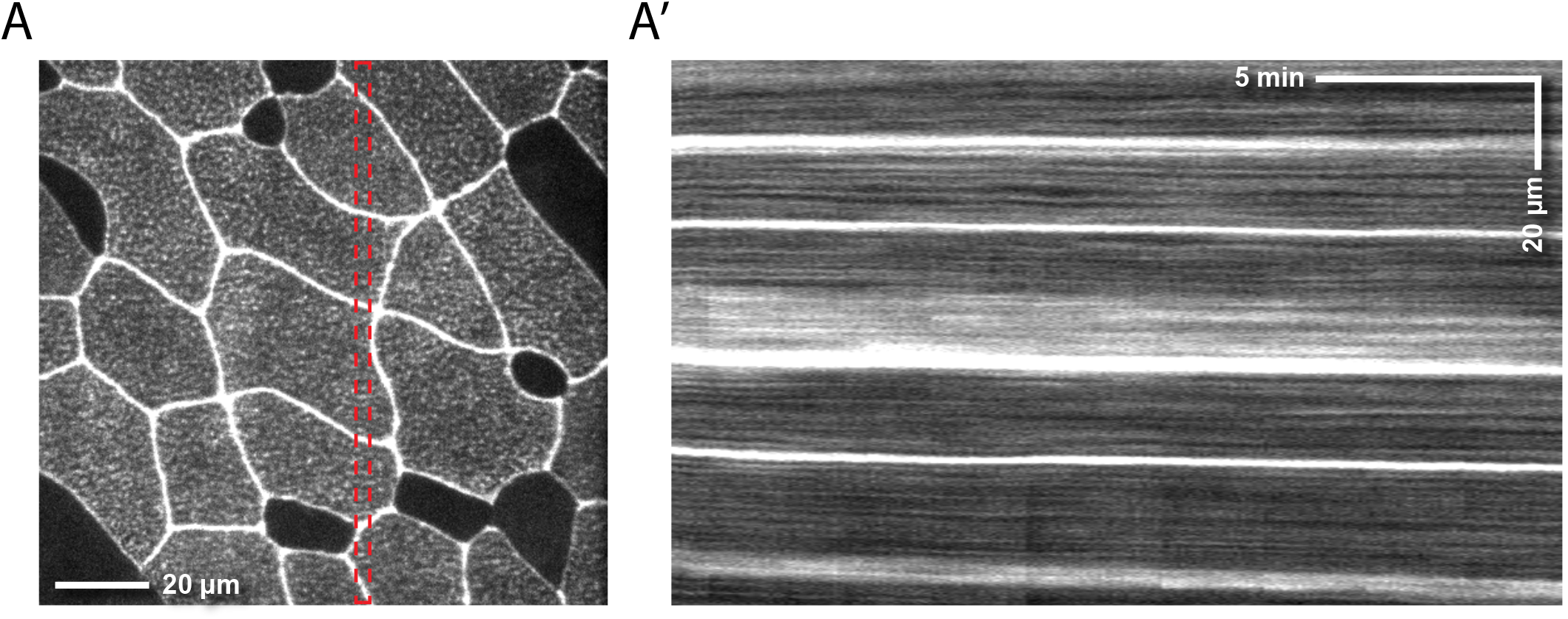
Lack of evidence for cortical excitability in stage 20 embryos. (A) Representative still image (left) of UtrCH labeling in a stage 20 embryo. Non-uniform labeling is due to non-uniform mRNA distribution of the GFP-UtrCH probe after microinjection. (A’) Kymograph of the dataset shown in A showing mostly stable F-actin distribution over time.

## Citations

Amiel A, Houliston E (2009). Three distinct RNA localization mechanisms contribute to oocyte polarity establishment in the cnidarian Clytia hemisphaerica. Dev Biol 327, 191–203.

Bastos RN, Penate X, Bates M, Hammond D, Barr FA (2012). CYK4 inhibits Rac1-dependent PAK1 and ARHGEF7 effector pathways during cytokinesis. J Cell Biol 198:865–880.

Bement WM, Leda M, Moe AM, Kita AM, Larson ME, Golding AE, Pfeuti C, Su KC, Miller AL, Goryachev AB, von Dassow G (2015). Activator-inhibitor coupling between Rho signaling and actin assembly make the cell cortex an excitable medium. Nat Cell Biol 17, 1471–1483.

Bischof J, Brand CA, Somogyi K, Májer I, Thome S, Mori M, Schwarz US, Lénárt P (2017). A cdk1 gradient guides surface contraction waves in oocytes. Nat Commun 8, 849.

Bischof J, Brand, CA, Somogyi K, Májer I, Thome S, Mori M, Schwarz US, Lénárt P (2017). A cdk1 gradient guides surface contraction waves in oocytes. Nature 8, 849.

Brownlee C, Heald R (2019). Importin α Partitioning to the Plasma Membrane Regulates Intracellular Scaling. Cell 176, 805–815.

Burkel BM, von Dassow G, Bement WM (2007). Versatile fluorescent probes for actin filaments based on the actin-binding domain of utrophin. Cell Motil Cytoskeleton 64, 822–832.

Canman JC, Lewellyn L, Laband K, Smerdon SJ, Desai A, Bowerman B, Oegema K (2008). Inhibition of Rac by the GAP activity of centralspindlin is essential for cytokinesis. Science 322:1543–1546.

Case LB, Waterman CM (2011). Adhesive F-actin waves: a novel integrin-mediated adhesion complex coupled to ventral actin polymerization. PLoS One 6, e26631.

Chanet S, Huynh JR (2020). Collective Cell Sorting Requires Contractile Cortical Waves in Germline Cells. Curr Biol 30, 4213–4226.

Chang JB, Ferrell JE Jr (2013). Mitotic trigger waves and the spatial coordination of the Xenopus cell cycle. Nature 500:603–7.

Chen P, Tomschik M, Nelson KM, Oakey J, Gatlin JC, Levy DL (2019). Nucleoplasmin is a limiting component in the scaling of nuclear size with cytoplasmic volume. J Cell Biol 218, 4063–4078.

Faire K, Waterman-Storer CM, Gruber D, Masson D, Salmon ED, Bulinski JC (1999). E-MAP-115 (ensconsin) associates dynamically with microtubules in vivo and is not a physiological modulator of microtubule dynamics. J Cell Sci 112, 4243–4255.

Gerhardt M, Ecke M, Walz M, Stengl A, Beta C, Gerisch G (2014). Actin and PIP3 waves in giant cells reveal the inherent length scale of an excited state. J Cell Sci 127, 4507–4517.

Gerhart J, Wu M, Kirschner M (1984). Cell Cycle Dynamics of an M-phase-specific Cytoplasmic Factor in Xenopus laevis Oocytes and Eggs. J Cell Biol 98, 1247–1255.

Good MC, Vahey MD, Skandarajah A, Fletcher DA, Heald R (2013). Cytoplasmic volume modulates spindle size during embryogenesis. Science 342, 856–860.

Goryachev AB, Leda M, Miller AL, von Dassow G, Bement WM (2016). How to make a static cytokinetic furrow out of traveling excitable waves. Small GTPases 7:65–70.

Graessl M, Koch J, Calderon A, Kamps D, Banerjee S, Mazel T, Schulze N, Jungkurth JK, Patwardhan R, Solouk D, Hampe N, Hoffmann B, Dehmelt L, Nalbant P (2017). An excitable Rho GTPase signaling network generates dynamic subcellular contraction patterns. J Cell Biol 216:4271–4285.

Hamaguchi MS, Hiramoto Y (1978). Protoplasmic movement during polar-body formation in starfish oocytes. Exp Cell Res 112, 55–62.

Hara K (1971). Cinematographic observation of “surface contraction waves” (SCW) during the early cleavage of axolotl eggs. Wilhelm Roux Arch Entwickl Mech Org 167, 183–186.

Hara K, Tydeman P (1979). Cinematographic Observation of an “Activation Wave” (AW) on the Locally Inseminated Egg of Xenopus laevis. Wilehm Roux Arch Dev Biol 186, 91–94.

Hara K, Tydeman P, Kirschner M (1980). A cytoplasmic clock with the same period as the division cycle in Xenopus eggs. Proc Natl Acad Sci 77, 462–466.

Hara Y, Merten CA (2015). Dynein-Based Accumulation of Membranes Regulates Nuclear Expansion in Xenopus laevis Egg Extracts. Dev Cell 33, 562–575.

Hazel J, Krutkramelis K, Mooney P, Tomschik M, Gerow K, Oakey J, Gatlin JC (2013). Changes in cytoplasmic volume are sufficient to drive spindle scaling. Science 342, 853–856

Hui KL, Kwak SI, Upadhyaya A (2014). Adhesion-dependent modulation of actin dynamics in Jurkat T cells. Cytoskeleton 71, 119–135.

Ishihara K, Nguyen PA, Wühr M, Groen AC, Field CM, Mitchison TJ (2014). Organization of early frog embryos by chemical waves emanating from centrosomes. Philos Trans R Soc Lond B Biol Sci 369:20130454.

Landino J, Leda M, Michaud A, Swider ZT, Prom M, Field C, Bement WM, Vecchiarelli AG, Goryachev AB, Miller AL (2021). Rho and F-actin self-organize within an artificial cell cortex. Curr Biol S0960-9822(21)01410-X.

Maître JL, Niwayama R, Turlier H, Nédélec F, Hiiragi T (2015). Pulsatile cell-autonomous contractility drives compaction in the mouse embryo. Nat Cell Biol 17, 849–855.

Miao Y, Bhattacharya S, Banerjee T, Abubaker-Sharif B, Long Y, Inoue T, Iglesias PA, Devreotes PN (2019). Wave patterns organize cellular protrusions and control cortical dynamics. Mol Syst Biol 15:e8585.

Michaud A, Swider ZT, Landino J, Leda M, Miller AL, von Dassow G, Goryachev AB, Bement WM (2021). Cortical excitability and cell division. Curr Biol 31, R553–R559.

Michaux JB, Robin FB, McFadden WM, Munro EM (2018). Excitable RhoA dynamics drive pulsed contractions in the early C. elegans embryo. J Cell Biol 217, 4230–4252.

Miller AL and Bement WM (2009). Regulation of cytokinesis by Rho GTPase flux. Nat Cell Biol 11, 71–77.

Miller KE, Brownlee C, Heald R (2020). The power of amphibians to elucidate mechanisms of size control and scaling. Exp Cell Res 392:112036.

Newport JW and Kirschner MW (1984). Regulation of the cell cycle during Xenopus development. Cell 37, 731–742.

Pal D, Ellis A, Sepúlveda-Ramírez SP, Salgado T, Terrazas I, Reyes G, De La Rosa R, Henson JH, Shuster CB (2020). Rac and Arp2/3-Nucleated Actin Networks Antagonize Rho During Mitotic and Meiotic Cleavages. Front Cell Dev Biol 8:591141.

Pérez-Mongiovi D, Chang P, Houliston E (1998). A propagated wave of MPF activation accompanies surface contraction waves at first mitosis in Xenopus. J Cell Sci 111, 385–393.

Rankin S and Kirschner MW (1997). The surface contraction waves of Xenopus eggs reflect the metachronous cell-cycle state of the cytoplasm. Curr Biol 7, 451–454.

Stankevicins L, Ecker N, Terriac E, Maiuri P, Schoppmeyer R, Vargas P, Lennon-Duménil AM, Piel M, Qu B, Hoth M, Kruse K, Lautenschläger F (2020). Deterministic actin waves as generators of cell polarization cues. Proc Natl Acad USA 117, 826–835.

Thévenaz P, Ruttimann UE, Unser M. (1998). A Pyramid Approach to Subpixel Registration Based on Intensity. IEEE Trans Image Process. 7:27–41.

Vicker MG (2000). Reaction-diffusion waves of actin filament polymerization/depolymerization in Dictyostelium pseudopodium extension and cell locomotion. Biophys Chem 84, 87–98.

Weiner OD, Marganski WA, Wu LF, Altschuler SJ, Kirschner MW (2007). An actin-based wave generator organizes cell motility. PLoS Biol 5, e221.

Xiao S, Tong C, Yang Y, Wu M (2017). Mitotic Cortical Waves Predict Future Division Sites by Encoding Positional and Size Information. Dev Cell 43, 493–506.

Zhuravlev Y, Hirsch SM, Jordan SN, Dumont J, Shirasu-Hiza M, Canman JC (2017). CYK-4 regulates Rac, but not Rho, during cytokinesis. Mol Biol Cell 28:1258–1270.

von Dassow G, Verbrugghe KJ, Miller AL, Sider JR, Bement WM (2009). Action at a distance during cytokinesis. J Cell Biol 187:831–45.

